# Neuromodulation of astrocytic K+ clearance

**DOI:** 10.1101/841643

**Authors:** Alba Bellot-Saez, Orsolya Kékesi, Yuval Ben-Abu, John W. Morley, Yossi Buskila

## Abstract

Potassium homeostasis is a fundamental requirement for normal brain function. Therefore, effective removal of excessive K^+^ accumulation from the synaptic cleft during neuronal activity is paramount. Astrocytes, one of the most common subtype of glial cells in the brain, play a key role in K^+^ clearance from the extracellular milieu using various mechanisms, including uptake via Kir channels and the Na^+^-K^+^ ATPase, and spatial buffering through the astrocytic gap-junction coupled network. Recently we showed that alterations in the concentrations of extracellular potassium ([K^+^]°) or impairments of the astrocytic clearance mechanism effect the resonance and oscillatory behaviour of both individual and networks of neurons recorded from C57/BL6 mice of both sexes. These results indicate that astrocytes have the potential to modulate neuronal network activity, however the cellular effectors that may affect the astrocytic K^+^ clearance process are still unknown. In this study, we have investigated the impact of neuromodulators, which are known to mediate changes in network oscillatory behaviour, on the astrocytic clearance process. Our results suggest that some neuromodulators (5-HT; NA) affect astrocytic spatial buffering via gap-junctions, while others (DA; Histamine) affect the uptake mechanism via Kir channels. These results suggest that neuromodulators can affect network oscillatory activity through parallel activation of both neurons and astrocytes, establishing a synergistic mechanism to recruitment of neurons into ensamble of networks to maximise the synchronous network activity.

**Significance statement:** Neuromodulators are known to mediate changes in network oscillatory behaviour and thus impact on brain states. In this study we show that certain neuromodulators directly affect distinct stages of astrocytic K^+^ clearance, promoting neuronal excitability and network oscillations through parallel activation of both neurons and astrocytes, thus establishing a synergistic mechanism to maximise the synchronous network activity.

## INTRODUCTION

Animal survival is highly dependent on their ability to adapt to the changing environment. To do so, animals are constantly switching between behavioral states, which are correlated to different network oscillations. We recently showed that local alterations in the extracellular K^+^ concentration ([K^+^]°) can affect the oscillatory activity of neuronal networks (Buskila *et al.*, 2019) and that specific impairments of the astrocytic clearance mechanism can affect the resonance and oscillatory behaviour of neurons both at single cell and network levels (Bellot-Saez *et al.*, 2018), implying that astrocytes can modulate neuronal network activity. However, the cellular and molecular mechanisms that affect this clearance process by astrocytes are still unknown.

Historically, network oscillations were considered to be highly affected by neuromodulation(Lee & Dan, 2012), and previous studies indicated an essential role for neuromodulators in mediating the shift between different behavioural states. However, we still know little about the circuitry involved in this neuromodulation, specifically at the cellular level.

Cortical astrocytes found to express a wide variety of receptors for different neuromodulators, including Acetylcholine (Ach, nicotinic α/β and metabotropic M^1-4^)(Oikawa *et al.*, 2005; Amar *et al.*, 2010), Histamine (H^1-3^)(N. Inagaki, H. Fukui, Y. Taguchi, N.P. Wang, A. Yamatodani, 1989), Serotonin (5-HT^1,2,5,6,7^)(Hirst *et al.*, 1998), Noradrenaline (NA, α_1,2_-adrenoreceptors and Β_1,2_-adrenoreceptors)(Morin *et al.*, 1996; Bekar *et al.*, 2008) and Dopamine (DA, D_1-5_)(Khan *et al.*, 2001; Wei *et al.*, 2018), which evoke [Ca^2+^]^i^ increases that affect astrocytic function. For instance, Histamine leads to astrocytic [Ca^2+^]^i^ increases *in vitro* (Jung *et al.*, 2000) and mediates the upregulation of the glutamate transporter 1 (GLT-1) through astrocytic H_1_ receptors, leading to reduced extracellular glutamate levels (Fang *et al.*, 2014) and thus playing a neuroprotective role against excitotoxicity. Similar to Histamine, NA (Ding *et al.*, 2013; Schnell *et al.*, 2016; Nuriya *et al.*, 2017), DA (Vaarmann *et al.*, 2010; Bosson *et al.*, 2015; Jennings *et al.*, 2017), 5-HT (Blomstrand *et al.*, 1999; Sandén *et al.*, 2000; Schipke *et al.*, 2011) and ACh (Araque *et al.*, 2002; Oikawa *et al.*, 2005; Perea, 2005) also elicit [Ca^2+^]^i^ elevations in astrocytes, and a recent study showed that superfusion of a cocktail containing these neuromodulators triggered a transient increase in the [K^+^]° levels (Ding *et al.*, 2016).

Astrocytic Ca^2+^ signalling and glutamate clearance play an essential role in the regulation of the network activity and K^+^ homeostasis, which ultimately affects neuronal excitability underlying network oscillations (Wang Fushun *et al.*, 2012; Ding *et al.*, 2016). Recently, Ma et al. (2016) (Ma *et al.*, 2016) showed that neuromodulators can signal through astrocytes by affecting their Ca^2+^ oscillations to alter neuronal circuitry and consequently behavioural output. In line with these observations, Nedergaard’s group further demonstrated that bath application of neuromodulators to cortical brain slices increased [K^+^]° regardless of synaptic activity (Ding *et al.*, 2016), suggesting that increased [K^+^]° could serve as a mechanism to maximize the impact of neuromodulators on synchronous activity and recruitment of neurons into networks.

In this study, we have investigated (i) which neuromodulators can affect astrocytic K^+^ clearance mechanisms (including K^+^ uptake through K_ir_4.1 channels and K^+^ spatial buffering via gap junctions (GJs) to amend [K^+^]° levels; and ii) does the impact of the different neuromodulators depend on [K^+^]° and thereby different levels of network activity?

To this end, we assessed the impact of specific neuromodulators on the astrocytic K^+^ clearance mechanism by measuring their influence on the [K^+^]° clearance time-course in acute brain slices and further assessed whether this impact was due to a direct activation of astrocytic receptors or due to indirect influence via the neural network.

## MATERIALS AND METHODS

### Animals and slice preparation

For this study, we used 4-8-week-old C57/BL6 mice of either sex. All animals were healthy and handled with standard conditions of temperature, humidity, twelve hours light/dark cycle, free access to food and water, and without any intended stress stimuli. All experiments were approved and performed in accordance with Western Sydney University committee for animal use and care guidelines (Animal Research Authority #A10588).

For slice preparations, animals were deeply anesthetized by inhalation of isoflurane (5%), decapitated, and their brains were quickly removed and placed into ice-cold physiological solution (artificial CSF, aCSF) containing (in mM): 125 NaCl, 2.5 KCl, 1 MgCl_2_, 1.25 NaH_2_PO_4_, 2 CaCl_2_, 25 NaHCO_3_, 25 glucose and saturated with carbogen (95% O_2_-5% CO_2_ mixture; pH 7.4). Parasagittal brain slices (300 μm thick) were cut with a vibrating microtome (Leica VT1200S) and transferred to the Braincubator™ (PaYo Scientific, Sydney), as reported previously (Buskila *et al.*, 2013). The Braincubator is an incubation system that closely monitors and controls pH, carbogen flow and temperature, as well as irradiating bacteria through a separate UV chamber (Breen & Buskila, 2014; Cameron *et al.*, 2016). Slices were initially incubated for 12 min at 35 °C, after which they were allowed to cool to 15–16 °C and kept in the Braincubator™ for at least 30 min before any measurement (Buskila *et al.*, 2014, 2020).

### Electrophysiological recording and stimulation

The recording chamber was mounted on an Olympus BX-51 microscope equipped with IR/DIC optics and Polygon 400 patterned illuminator (Mightex). Following staining (Fluo-4 AM, SR101) and short recovery period in the Braincubator™, slices of somatosensory cortex were mounted in the recording chamber, for a minimum of 15 minutes, to allow them to warm up to room temperature (∼22°C) and were constantly perfused at a rate of 2 ml/min with carbogenated aCSF.

[K^+^]° measurements were performed in layer II/III of the somatosensory cortex, by placing the K^+^-selective microelectrode nearby a selected astrocyte (termed “astrocyte ∝”) stained with SR101 (Figure 1A). K^+^ clearance is temperature-dependent, with Q^10^ of 1.7 at 26 °C and 2.6 at 37°C, mainly due to Na^+^/K^+^ ATPase activity (Ransom *et al.*, 2000*a*). Due to the absence of a selective blocker for astrocytic Na^+^/K^+^ ATPase, and our interest in assessing K^+^ uptake into astrocytes via Kir channels, these measurements were performed at room temperature (22°C), when Na^+^/K^+^ ATPase activity is fairly low. Various KCl concentrations, corresponding to low (∼5 mM), high (∼15 mM) and excessive (∼30 mM), were added to physiological aCSF and locally applied at a constant distance (∼10 µm) from the K^+^-selective microelectrode through a puffing pipette (tip diameter of 1 μm) for 0.1 sec, as previously described (Bellot-Saez *et al.*, 2018). Preparation and calibration of the K^+^-selective microelectrodes were performed as detailed in (Deveau *et al.*, 2005; Haack & Rose, 2014). In short, the voltage response of the silanized K^+^-selective microelectrodes was calibrated before and after experiments within the experimental chamber by placing the electrode in aCSF containing different KCl concentrations (2.5 or “normal” aCSF, 4, 10, 15 or 30 mM). Once the electrode potential reached a steady state, a dose-response curve was calculated using a half-logarithmic (Log^10^) plot. K^+^-selective microelectrodes were considered good if the recorded voltage baseline was stable and the voltage response was similar before and after its experimental usage (∼10 % deviation). The K^+^ clearance rate was calculated by dividing the concentration amplitude with the decay time (90-10%).

**Figure 1.**
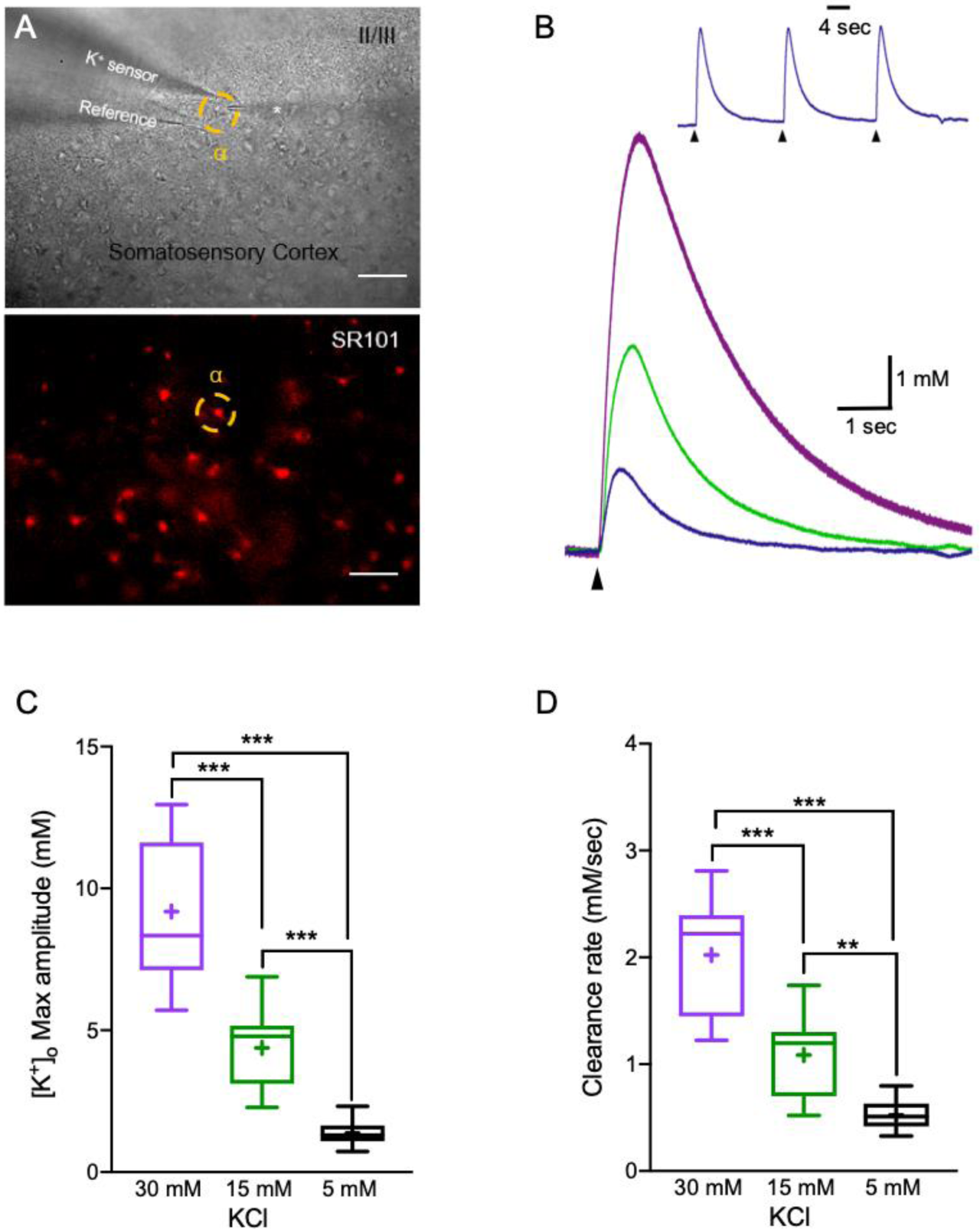
Measuring K^+^ clearance in acute brain slices. A) Top - DIC image of a brain slice depicting the the K^+^ recording electrode together with the KCl puffing electrode in layers 2/3 of the somatosensory cortex (scale bar - 50 μm). Bottom – fluorescence image of the same region depicting astrocytes stained with SR101; yellow circle surrounds to an astrocyte nearby the recording electrode (astrocyte ∝). B) Top-sample traces of [K^+^]° recordings following repetitive stimulation with 5 mM KCl, arrows indicate the time of KCl application. Bottom-average traces of 3 repetitive [K^+^]° recordings depicting the K^+^ clearance time course following local application of 30 mM (purple), 15 mM (green) and 5 mM (blue) KCl puffs. C,D) Box plots depicting the maximal amplitude (C) and clearance rate (D) following local application of 5, 15, and 30 mM KCl. The box upper and lower limits are the 25^th^ and 75^th^ quartiles, respectively. The whiskers depicting the lowest and highest data points, while the + sign represents the mean and the horizontal line through the box is the median.

**Figure 2.**
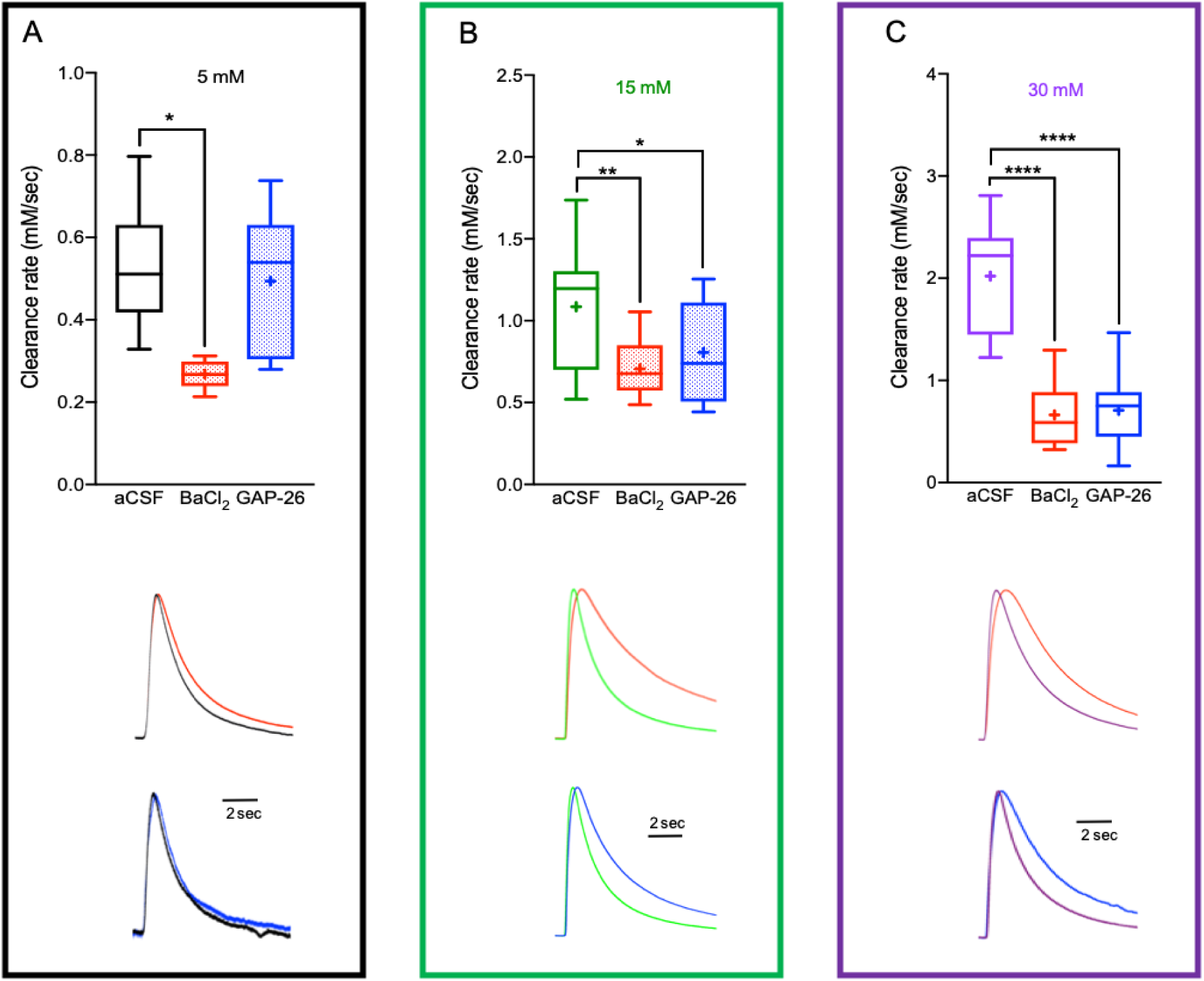
Impaired astrocytic K^+^ clearance affect [K^+^]° dynamics. A-C) Box plots depicting the average K^+^ clearance rate following local application of 5 mM (A) 15 mM (B), and 30 (C) mM KCl, under control conditions (aCSF), reduced Kir4.1 activity (red) or impaired astrocytic GJ conductance (blue). Bottom– sample traces of [K^+^]° recording (normalised to the peak amplitude) under control and impaired clearance conditions; red traces-with BaCl_2_; blue traces-with Gap-26/27 mixture. Control traces are colour coded as per their concentration (black-5 mM, green-15 mM and purple-30 mM). Box plots definition are the same as in Fig. 1. *p < 0.05; **p < 0.01; ***p < 0.0001; two-way ANOVA.

To assess the impact of neuromodulators on the K^+^ clearance rate, [K^+^]° measurements were performed within the same brain slices before and after 5-minute bath application of different neuromodulators, including the cholinergic agonist Carbachol (100 μM), Histamine dihydrochloride (50 μM), Noradrenaline bitartrate (40 μM), NPEC-caged-Serotonin (30 μM) and NPEC-caged-Dopamine (10 μM). To exclude the involvement of neuronal activity, similar experiments were conducted after perfusing slices with neuromodulators and tetrodoxin (TTX, 1 μM) for 5 additional minutes. Polygon400 illuminator (Mightex) was used to uncage NPEC-caged-Serotonin and NPEC-caged-Dopamine compounds by applying focal photolysis with UV light (∼360 nm) in a selected area (50 µm) which includes the surroundings of the K^+^-selective microelectrode, the KCl puffing pipette and the selected astrocytic domain with its processes, for 1 second prior to local application of KCl (Figure 3A).

**Figure 3.**
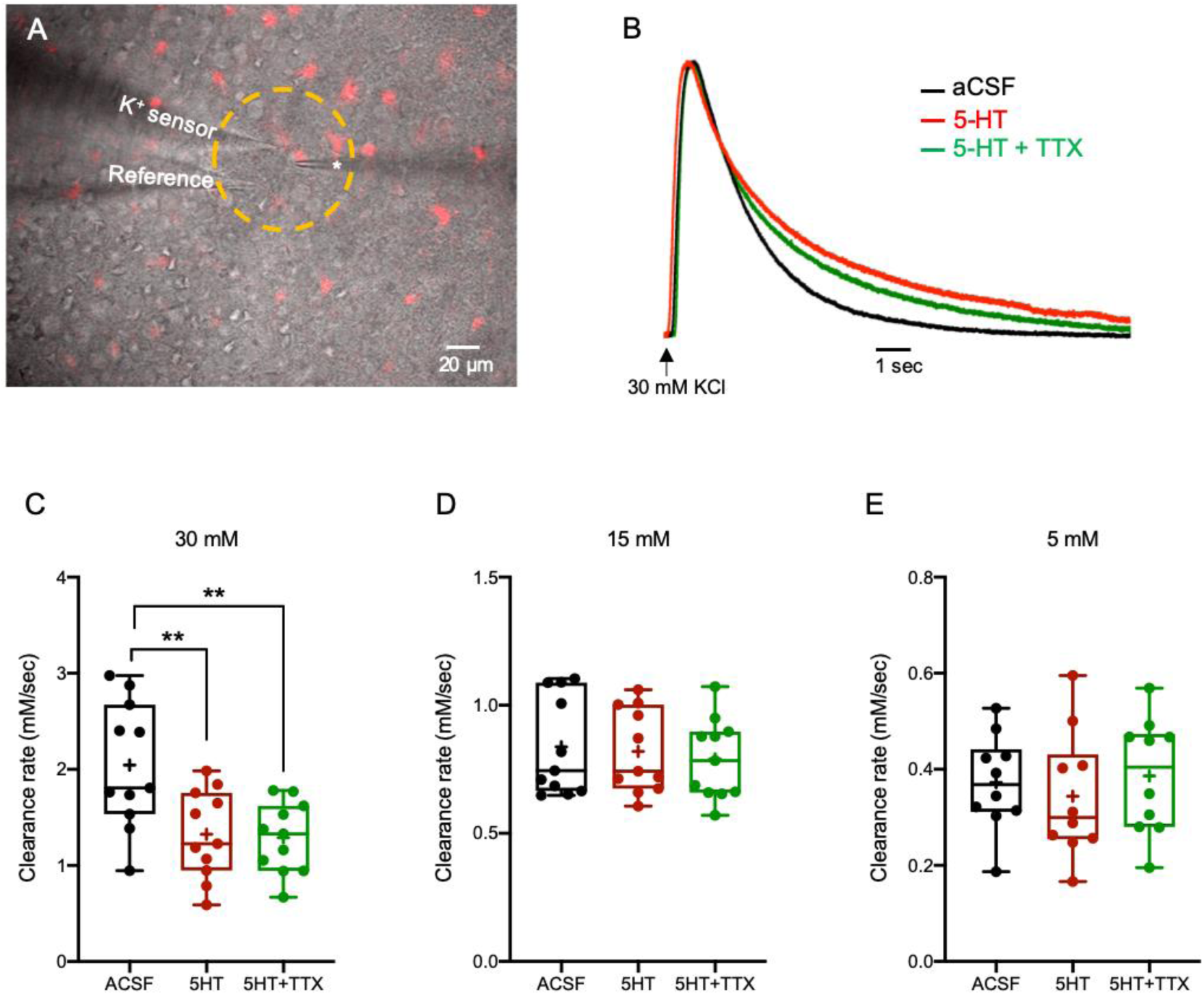
The impact of 5-HT on the K^+^ clearance rate and astrocytic Ca^2+^ signalling. A) Merged image of DIC (grey) and SR-101 (red) staining depicting the experimental setup, including the K^+^ recording electrode, the KCl puffing electrode (*) and the photo-activated region (yellow circle, 360 nm for 1 second prior to local application of KCl) used for application of the neuromodulators Serotonin (NPEC-caged-Serotonin, 30 μM) and Dopamine (NPEC-caged-Dopamine, 10 μM). B) Sample traces of [K^+^]° recordings depicting the K^+^ clearance time course following local application of 30 mM KCl (arrow), before (aCSF, black) and after focal photolysis of 30 μM caged Serotonin (5-HT, red) or 30 μM caged 5-HT with 1 μM TTX (green). C-E) Box plots depicting the K^+^ clearance rate following local application of 30 mM (C), 15 mM (D) and 5 mM (E) KCl before (black dots) and after photolysis of 5-HT (red dots) or 5-HT + TTX (green dots). Box plot definitions is the same as in Fig. 1. *\*p < 0.05; **p < 0.01; ***p < 0.0001*

### Drugs

All drugs were stored as frozen stock solutions and were added to aCSF just before recordings. Neuromodulators, including NA, Histamine, 5-HT and DA were purchased from Tocris Bioscience (In vitro Technologies Pty Ltd). Noradrenaline bitartrate and Histamine dihydrochloride were dissolved in water to a stock solution at final concentration of 100 mM. Carbachol (Sigma Aldrich) and caged neuromodulators, including NPEC-caged-Serotonin and NPEC-caged-Dopamine, were dissolved in DMSO to a stock solution at final concentrations of 1 M or 100 mM, respectively. All stock solutions were stored at −°C and protected from light when required.

### Experimental Design and Statistical analysis

Detailed experimental designs of [K^+^]° measurement and Ca^2+^ signalling studies are described in Materials and Methods, and Results, including the number of animals used and cells included in the analyses. These numbers were based on our previous studies and standard practices in the field.

Comparrisons of K^+^ clearance before and after application of a certain neuromodulator or TTX were conducted using two-tailed paired student t-test, as they were conducted on same slices and in the same region. For group comparrisons between different [K^+^]° concentrations or treatments, in which different slices from different animals were used, we conducted one-way or two-way ANOVA followed by Tukey’s post hoc test, according to the experimental design. Statistical comparisons were done with Prism 7 (GraphPad Software; San Diego, CA), and unless stated, data is reported as mean ± S.E.M. Analysis of K^+^ transient properties were performed using a custom-made MATLAB code (MathWorks). Probability values < 0.05 were considered statistically significant.

## RESULTS

### K^+^ clearance time course in acute brain slices

To measure the [K^+^]° clearance rate, we have used custom built double-barreled K^+^-selective microelectrodes, as previously described (Haack *et al.*, 2015). Transient elevations of [K^+^]° were mediated via local application of KCl at various concentrations corresponding to low (5 mM), high (15 mM) and excessive (30 mM) [K^+^]°, which correlate to different levels of network activity. K^+^-selective microelectrodes were placed in layer II/III of the somatosensory cortex, (close to a selected astrocyte stained with SR101 named “astrocyte ∝”, Figure 1A) and were calibrated before and after each experiment, as described in Materials & Methods.

Under control aCSF conditions, local application of 30 mM, 15 mM or 5 mM KCl led to [K^+^]° increases of 9.19±0.65 mM (n=14), 4.38±0.34 (n=16) and 1.38±0.11 (n=15) respectively (F(2, 42) = 88.15, p < 0.0001, one-way ANOVA with Tukey’s post hoc test, Figure 1 B, C). The differences between the applied KCl concentrations and the measured [K^+^]° concentrations (by the K^+^ selective electrode located 10 µm away from astrocyte ∝) were due to the dilution of the applied KCl solution.

The [K^+^]° clearance rate was defined as the time it took the [K^+^]° concentration to return to baseline levels measured before the local application of external KCl (90-10% decreasing slope, Figure 1B, D). Under control aCSF conditions, the [K^+^]° clearance rate was concentration-dependent, ranging from 2.02±0.14 mM/sec following local application of 30 mM KCl (n=14) and decreased to 1.09±0.09 mM/sec (n=16) and 0.56±0.05 mM/sec (n=15) following 15 and 5 mM KCl respectively (F(2, 46) = 62.03, p < 0.0001, one-way ANOVA with tukey’s post hoc test, Figure 1D). These results are consistant with previous study showing that the decay rate of [K^+^]° was inversly correlated to the [K^+^]° amplitude (Ransom *et al.*, 2000*a*).

### Alterations in astrocytic K^+^ uptake and buffering mechanisms affect the [K^+^]° clearance rate

We next assessed the specific impact of astrocytic K^+^ clearance mechanisms, including “uptake via Kir4.1” and “spatial buffering via GJ”, on the [K^+^]° clearance rate. Bath application of BaCl_2_ (100 µM) that previously shown to selectively block astrocytic K_ir_4.1 channels and thus the uptake into astrocytes (Larsen & Macaulay, 2014; Ma *et al.*, 2014), significantly (*F*_(_1, 83) = 103.6, p < 0.0001, two-way ANOVA) reduced the K^+^ clearance rate for all [K^+^]° tested (30 mM, 0.66±0.07 mM/sec, n=18; 15 mM, 0.71±0.04 mM/sec, n=14; 5 mM, 0.28±0.01 mM/sec, n=12; p < 0.0001, two-way ANOVA with Tukey’s post hoc test, Figure 2A-C), indicating a slower rate of K^+^ removal from the extracellular space when K^+^ uptake is impaired.

To selectively block K^+^ spatial buffering through the astrocytic network, we incubated the slices with a mixture of connexin-43 (Cx43) mimetic peptides (GAP-26, 200μM and GAP-27, 300μM), that selectively decrease astrocytic connectivity via electrical GJs (Bellot-Saez *et al.*, 2018). Indeed, selective blockade of Cx43 significantly reduced the K^+^ clearance rate (F(1, 82) = 58.54, p < 0.0001, two-way ANOVA). However, disruption of the astrocytic connectivity had a differential impact on the K^+^ clearance rate, as it affected K^+^ transients only at high (15 mM, 0.81±0.04 mM/sec, n=15, Figure 2B) and excessive (30 mM, 0.71±0.08 mM/sec, n=17, Figure 2C) [K^+^]° levels (p < 0.0001, two-way ANOVA with tukey’s post hoc test). Low [K^+^]° did not lead to a significant change in the K^+^ clearance rate (5 mM, 0.50±0.03 mM/sec, n=11; p = 0.84, two-way ANOVA with tukey’s post hoc test, figure 2A), confirming the hypothesis that K^+^ uptake via the Na^+^-K^+^ ATPase and Kir4.1 channels is the dominant process used to clear low levels of [K^+^]° and astrocytic K^+^ spatial buffering via GJ takes place at higher levels of network activity (Enkvist & McCarthy, 1994). These results are also in agreement with a computational model constructed to simulate the impact of “net uptake” and spatial buffering on [K^+^]° dynamics (Halnes *et al.*, 2013).

### Neuromodulation of astrocytic K^+^ clearance; Serotonin (5-HT)

Previous studies have demonstrated that astrocytes express several subtypes of serotonergic receptors across different brain areas, including the cortex, corpus callosum, brain stem, spinal cord and hippocampus (Bonhaus *et al.*, 1995; Azmitia *et al.*, 1996; Carson *et al.*, 1996; Hirst *et al.*, 1998; Sandén *et al.*, 2000; Maxishima *et al.*, 2001; Mahé *et al.*, 2004). Cortical astrocytes have been found to express 5HT_2b_ receptors coupled to phospholipase A_2_ (PLA_2_) and phospholipase C (PLC)/G_q_ signalling cascades, whose activation leads to Ca^2+^ release from internal stores (Li *et al.*, 2010) and stimulation of glycogenolysis (Kong *et al.*, 2002). To test the impact of 5-HT on the K^+^ clearance rate, we photolyzed (50 µm diameter; UV light) NPEC-caged-Serotonin (30 µM) in layer II/III of the somatosensory cortex, which include the astrocytic domain and nearby neurons (Figure 3A).

Our results show that following transient application of excessive KCl concentration (30 mM, n=11) and uncaging of 5-HT, the K^+^ clearance rate decreased to 1.33±0.14 mM/sec (F(1.295, 12.95) = 20.68, p = 0.0003, one-way ANOVA with Tukey’s post hoc test, Figure 3B, C). However, co-application of 5-HT and lower KCl concentrations (15 mM and 5 mM) did not affect the K^+^ clearance rate significantly (0.82±0.05 mM/sec, n=11 and 0.34±0.04 mM/sec, n=10 respectively; (F(1.420, 14.20) = 2.549, p = 0.1246, one-way ANOVA, Figure 3D, E, Supplementary table 1). Moreover, blockade of neuronal spiking activity with TTX (1 µM) prior to 5-HT photolysis was comparable with the effect of 5-HT alone, suggesting that the observed alterations in the K^+^ clearance rate at excessive [K^+^]° (∼30 mM) are independent of neuronal activity and likely due to the direct effect of 5-HT on the astrocytic spatial buffering mechanism.

**Table 1.**
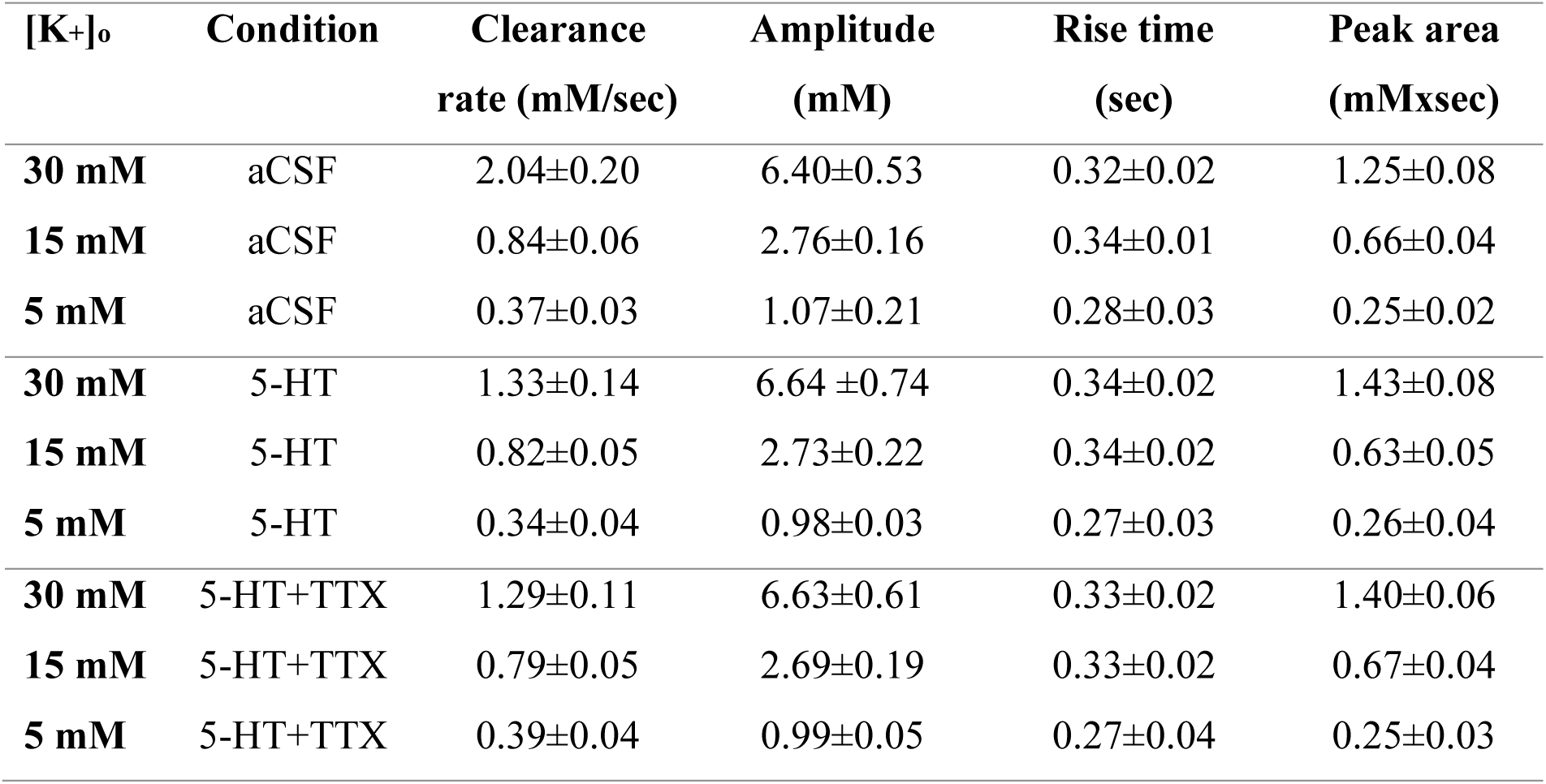
The impact of 5-HT on the K^+^ clearance rate. Average values of the [K^+^]° clearance rate (90-10%), amplitude, rise time (10-90%) and top peak area (10%) at all concentrations tested, before and after the application of 5-HT or 5-HT + TTX. Data is reported as mean ± S.E.M.

### Dopamine (DA)

DA receptors are classically grouped into D_1_-like (D_1_ and D_5_) and D_2_-like (D_2_, D_3_ and D_4_) receptors that activate opposite signalling cascades(Wei *et al.*, 2018). While DA binding to D_1_-like receptors promotes an increase in 3’,5’-cyclic adenosine monophosphate (cAMP) levels and the activation of protein kinase A (PKA) via adenylyl cyclase (Ac)(Zanassi *et al.*, 1999), activation of D_2_-like receptors, which are coupled to PLC/inositol 1,4,5-triphosphate (IP_3_) pathway, triggers Ca^2+^ release from internal stores and decreases cAMP levels (Khan *et al.*, 2001).

In order to assess the overall impact of DA on the K^+^ clearance rate we locally uncaged NPEC-caged-Dopamine compounds (10 µM)(Castro *et al.*, 2013), as described above for 5-HT (Figure 3A). Focal photolysis of caged DA significantly reduced the K^+^ clearance rate at all [K^+^]° concentrations tested, including 30 mM (1.68±0.25 mM/sec, n=13, (F(1.579, 18.95) = 108.5), p<0.0001, one-way ANOVA with Tukey’s post hoc test), 15 mM (1.21±0.15 mM/sec, n=12, (F(2, 22) = 6.886, p = 0.0048, one-way ANOVA with Tukey’s post hoc test) and 5 mM (0.56±0.08 mM/sec, n=14, (F(1.58, 20.54) = 15.15, p = 0.0002, one-way ANOVA with Tukey’s post hoc test, Figure 4A-D, Supplementary table 2). Moreover, this effect was independent of neuronal activity, as addition of TTX (1 µM) displayed the same results (Figure 4A-D).

**Table 2.** The impact of DA on the K^+^ clearance rate. Average values of the [K^+^]° clearance rate (90-10%), amplitude, rise time (10-90%) and top peak area (10%) at all concentrations tested, before and after the application of DA or DA + TTX. Data is reported as mean ± S.E.M.

| [K<sup>+</sup>]<sub>o</sub> | Condition | Clearance rate<br>(mM/sec) | Amplitude<br>(mM) | Rise time<br>(sec) | Peak area<br>(mMxsec) |
| --- | --- | --- | --- | --- | --- |
| **30 mM** | aCSF | 2.46±0.28 | 7.51±1.05 | 0.27±0.03 | 1.96±0.15 |
| **15 mM** | aCSF | 1.60±0.25 | 4.24±0.43 | 0.27±0.02 | 0.99±0.08 |
| **5 mM** | aCSF | 0.80±0.11 | 1.46±0.16 | 0.26±0.02 | 0.35±0.02 |
| **30 mM** | DA | 1.61±0.26 | 7.32±0.83 | 0.28±0.03 | 2.41±0.17 |
| **15 mM** | DA | 1.35±0.17 | 4.13±0.40 | 0.29±0.02 | 1.21±0.11 |
| **5 mM** | DA | 0.60±0.09 | 1.49±0.19 | 0.27±0.02 | 0.46±0.04 |
| **30 mM** | DA+TTX | 1.68±0.25 | 7.25±0.82 | 0.28±0.02 | 2.36±0.22 |
| **15 mM** | DA+TTX | 1.21±0.15 | 4.26±0.41 | 0.28±0.01 | 1.10±0.09 |
| **5 mM** | DA+TTX | 0.56±0.08 | 1.49±0.16 | 0.27±0.01 | 0.42±0.04 |

**Figure 4.**
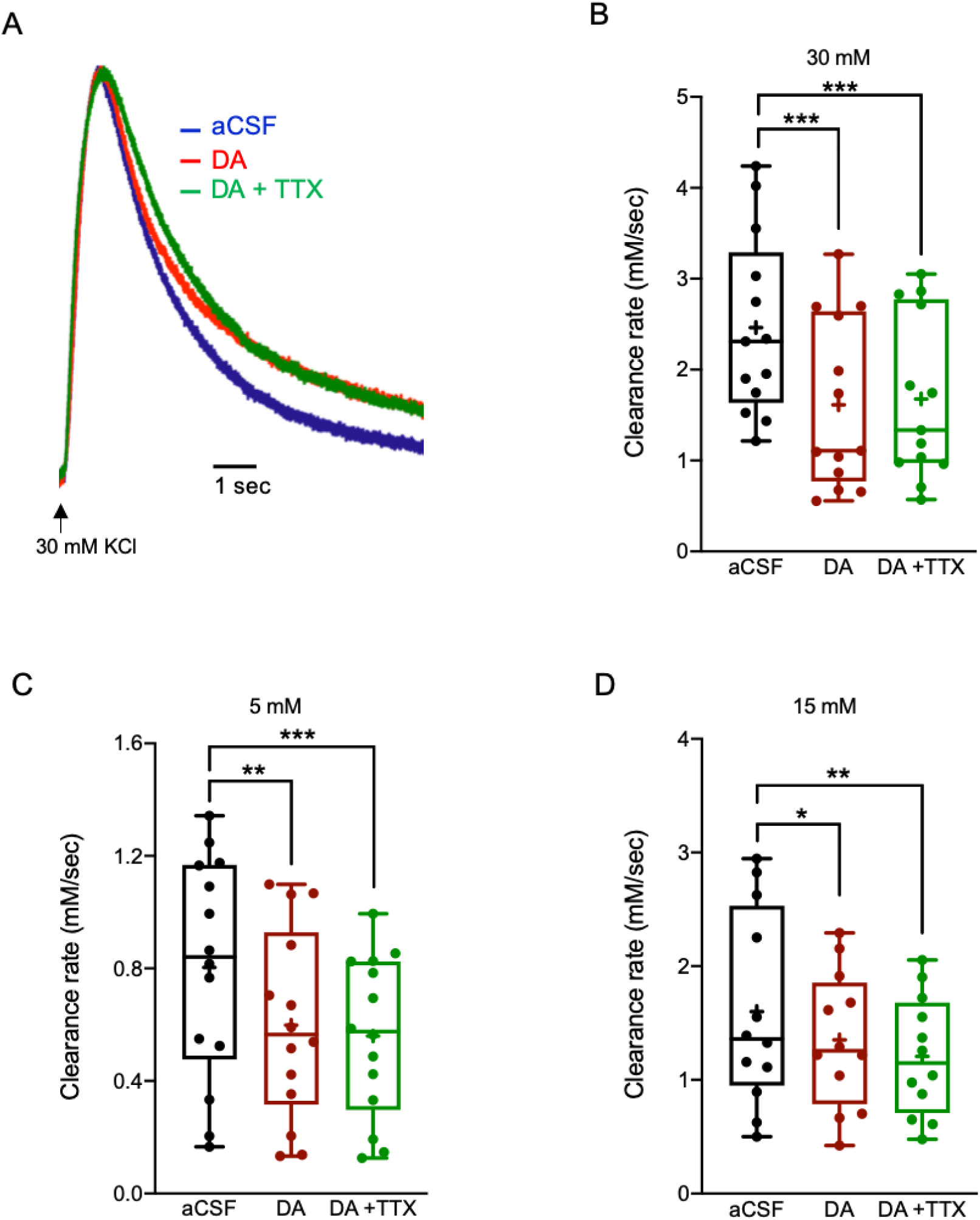
The impact of DA on the K^+^ clearance rate and astrocytic Ca^2+^ signalling. A) Sample traces of [K^+^]° recordings depicting the K^+^ clearance time course following local application of 30 mM KCl (arrow), before (aCSF, black) and after focal photolysis of 30 μM caged Dopamine (DA, red) or 30 μM caged DA with 1 μM TTX (green). B-D) Box plots depicting the K^+^ clearance rate following local application of KCl at 30 mM (B), 15 mM (C) and 5 mM (D), before (aCSF, black dots) and after DA uncaging without (red dots) or with TTX (green dots). Box plot definitions is the same as in Fig. 1. *p < 0.05; **p < 0.01; ***p < 0.0001

### Noradrenaline (NA)

Astrocytes express several receptors for NA, including α_1,_ α_2_ and β_1_-adenergic receptors, which mediate multiple processes. Activation of α_1_ receptors triggers the PLC/IP_3_ signalling cascade that results in Ca^2+^ release from the internal stores (Duffy & MacVicar, 1995; Ding *et al.*, 2013) leading to enhanced activity of protein kinase C (PKC) and the cAMP response element-binding (CREB)-dependent transcription (Carriba *et al.*, 2012), and also exacerbates glutamate re-uptake into astrocytes through GLT-1/GLAST glutamate transporters (Alexander *et al.*, 1997). In contrast, activation of α_2_ receptors in astrocytes primarly increase glycogenesis and reduces cAMP activity via the inhibitory G-protein (Gi/o), thus providing high ATP levels during periods of high demand, although under certain conditions it may permit glycogenolysis (Donnell *et al.*, 2012). However, stimulation of astrocytic β1-adenergic receptors activates G-proteins (G_s_) which results in [Ca^2+^]^i^ increases (Nuriya *et al.*, 2017), cAMP accumulation, PKA activation and glycogenolysis (Xu *et al.*, 2014). Moreover, activation of β1-adenergic receptors enhance the Na^+^-K^+^-ATPase activity and thus facilitates K^+^ clearance following high neuronal activity, yet this effect is abolished at high non-physiological levels of [K^+^]° (Hajek *et al.*, 1996*b*).

Our results show that bath application of Noradrenaline bitartrate (40 µM) led to a decrease of the K^+^ clearance rate following local application of excessive (30 mM, 0.80±0.06 mM/sec, n=15, (F(1.485, 20.80) = 21.12), p < 0.0001, one-way ANOVA with Tukey’s post hoc test) or high [K^+^]° (15 mM, 0.70±0.06 mM/sec, n=16, F(1.396, 20.95) = 23.88, p < 0.0001, one-way ANOVA with Tukey’s post hoc test) regardless of neuronal activity, as the addition of TTX did not reverse this effect (Figure 5A-C). However, NA did not affect the K^+^ clearance rate at low [K^+^]° (5 mM, 0.42±0.04 mM/sec, n=15; (F(1.495, 20.92) = 1.423, p=0.2586, one-way ANOVA with Tukey’s post hoc test, Figure 5D, Supplementary table 3), suggesting it mainly affects the K^+^ spatial buffering process via gap junctions activated at high [K^+^]° and to less extent the uptake mechanism through Kir4.1 channels and Na/KATPase.

**Table 3.** The impact of NA on the K^+^ clearance rate. Average values of the [K^+^]° clearance rate (90-10%), amplitude, rise time (10-90%) and top peak area (10%) at all concentrations tested, before and after the application of NA or NA + TTX. Data is reported as mean ± S.E.M.

| [K<sup>+</sup>]<sub>o</sub> | Condition | Clearance rate<br>(mM/sec) | Amplitude<br>(mM) | Rise time<br>(sec) | Peak area<br>(mMxsec) |
| --- | --- | --- | --- | --- | --- |
| **30 mM** | aCSF | 1.42±0.14 | 5.99±0.26 | 0.40±0.02 | 1.20±0.08 |
| **15 mM** | aCSF | 0.87±0.05 | 2.56±0.15 | 0.35±0.02 | 0.54±0.06 |
| **5 mM** | aCSF | 0.44±0.04 | 0.92±0.06 | 0.30±0.02 | 0.25±0.04 |
| **30 mM** | NA | 0.80±0.06 | 5.94±0.45 | 0.39±0.03 | 1.45±0.07 |
| **15 mM** | NA | 0.70±0.06 | 2.52±0.11 | 0.36±0.02 | 0.78±0.08 |
| **5 mM** | NA | 0.42±0.04 | 0.92±0.05 | 0.29±0.02 | 0.30±0.04 |
| **30 mM** | NA+TTX | 0.90±0.07 | 5.76±0.41 | 0.38±0.03 | 1.54±0.10 |
| **15 mM** | NA+TTX | 0.65±0.05 | 2.49±0.09 | 0.35±0.02 | 0.77±0.08 |
| **5 mM** | NA+TTX | 0.42±0.04 | 0.91±0.04 | 0.30±0.03 | 0.28±0.05 |

**Figure 5.**
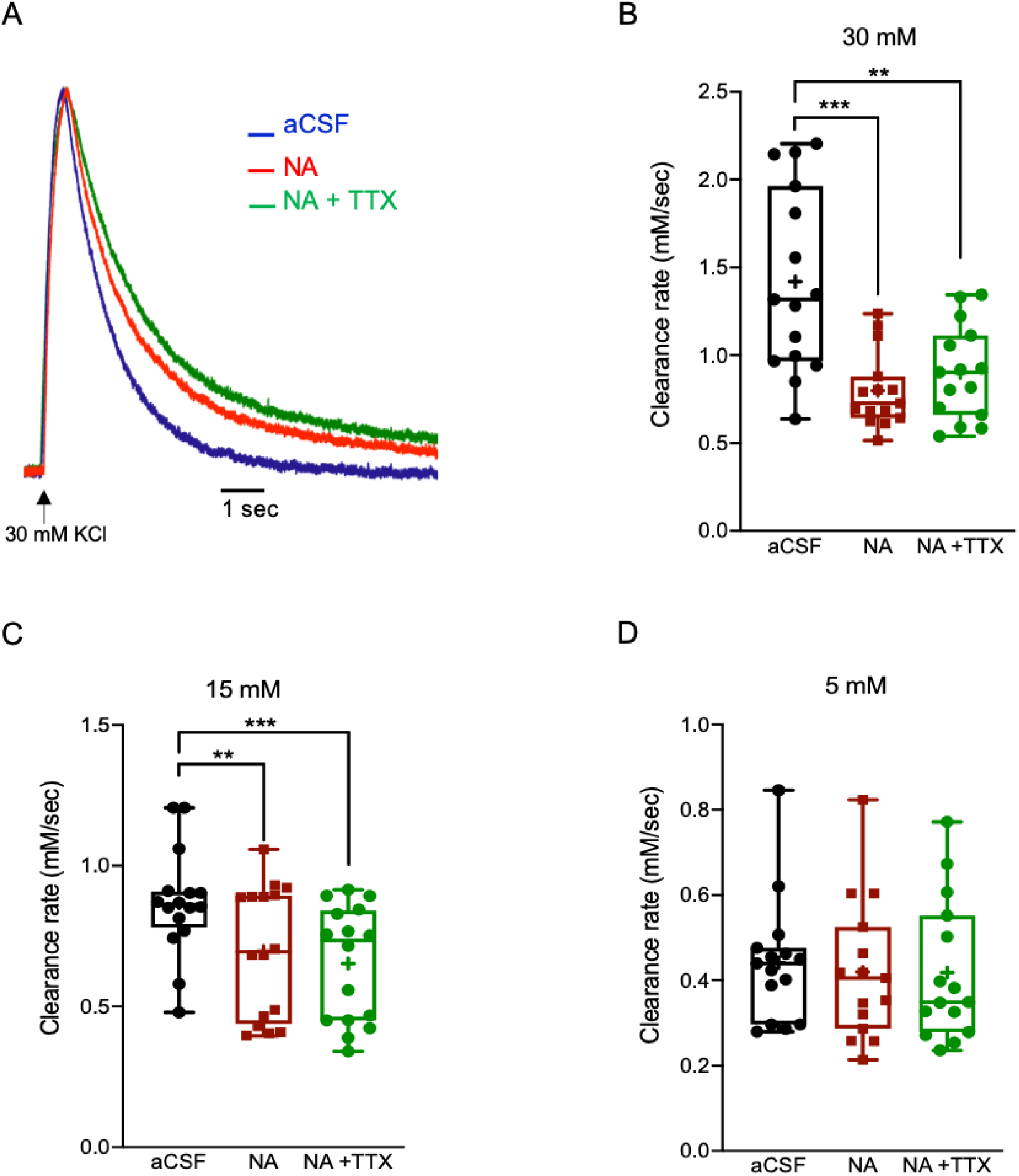
The impact of NA on the K^+^ clearance rate and astrocytic Ca^2+^ signalling. A) Sample traces of [K^+^]° recordings depicting the K^+^ clearance time course following local application of 30 mM KCl (arrow), before (aCSF, blue) and after application of Noradrenaline bitartrate (40 μM, NA, red) or NA with 1 μM TTX (green). B-D) Box plots depicting the K^+^ clearance rate following local application of KCl at 30 mM (B), 15 mM (C) and 5 mM (D), before (aCSF, black dots) and after bath application of NA without (red dots) or with TTX (green dots). Box plot definitions is the same as in Fig. 1. *p < 0.05; **p < 0.01; ***p < 0.0001

### Histamine

Astrocytes express different types of histaminergic receptors, including H_1_, H_2_ and H_3_, which mediate multiple processes, including glutamate clearance (Fang *et al.*, 2014) and glucose homeostasis (Medrano *et al.*, 1992). H_1_ receptors are G_q/11_-coupled and therefore associated with PKC and PLC signalling pathways, which lead to Ca^2+^ release from the endoplasmic reticulum (ER) (Arbonés *et al.*, 1988). H_2_ receptors are G_s_-coupled and have been found to participate in glycogen breakdown and energy supply via activation of PKA and stimulation of AC (Hill, 1990). H_3_ receptors are Gα_i/o_-coupled and less abundant in cortical astrocytes compared to astrocytes from other brain regions (e.g. striatum, hippocampus)(Mele & Jurič, 2013). These receptors have been involved in mediating the inhibition of AC, while triggering PLA_2_, MAP kinase and PI3K/AKT signalling pathways (Mariottini *et al.*, 2009; Jurič *et al.*, 2016).

Bath application of Histamine dihydrochloride (50 µM) significantly reduced the K^+^ clearance rate following local application of excessive, high and low KCl (30 mM, 1.15±0.14 mM/sec, n=10, F(1.013, 9.116) = 12.07, p = 0.0068, one-way ANOVA with Tukey’s post hoc test); 15 mM, 0.84±0.08 mM/sec, n=10, F(1.427, 12.84) = 16.60, p = 0.0006, one-way ANOVA with Tukey’s post hoc test); 5 mM, 0.30±0.02 mM/sec, n=11, F(1.566, 15.66) = 17.16, p = 0.0002, one-way ANOVA with Tukey’s post hoc test), Figure 6A-D). However, while the histaminergic impact on the K^+^ clearance rate at excessive [K^+^]° was not affected by neuronal activity (30 mM, 1.19±0.16 mM/sec, p= 0.22, one-way ANOVA with Tukey’s post hoc test, Figure 6B), blockade of neuronal firing with TTX increased the K^+^ clearance rate at high (15 mM, p= 0.01, one-way ANOVA with Tukey’s post hoc test, Figure 6C) and low [K^+^]° (5 mM, p= 0.04, one-way ANOVA with Tukey’s post hoc test, Figure 6D, Supplementary table 4), indicating the involvement of neuronal activity in mediating the histaminergic effects at these lower concentrations. These results suggest that the histaminergic regulation of astrocytic K^+^ clearance mechanisms is [K^+^]°-dependent and involves direct and indirect activation via the neural network.

**Table 4.** The impact of Histamine on the K^+^ clearance rate. Average values of the [K^+^]° clearance rate (90-10%), amplitude, rise time (10-90%) and top peak area (10%) at all concentrations tested, before and after the application of Histamine or Histamine + TTX. Data is reported as mean ± S.E.M.

| [K<sup>+</sup>]<sub>o</sub> | Condition | Clearance<br>rate (mM/sec) | Amplitude<br>(mM) | Rise time<br>(sec) | Peak area<br>(mMxsec) |
| --- | --- | --- | --- | --- | --- |
| **30 mM** | aCSF | 2.02±0.38 | 6.52±0.77 | 0.31±0.01 | 0.74±0.03 |
| **15 mM** | aCSF | 1.12±0.09 | 3.50±0.32 | 0.28±0.01 | 0.51±0.04 |
| **5 mM** | aCSF | 0.51±0.05 | 0.93±0.09 | 0.26±0.01 | 0.15±0.01 |
| **30 mM** | Histamine | 1.15±0.14 | 6.63±0.57 | 0.32±0.01 | 1.04±0.06 |
| **15 mM** | Histamine | 0.84±0.08 | 3.49±0.56 | 0.27±0.01 | 0.72±0.04 |
| **5 mM** | Histamine | 0.30±0.02 | 0.93±0.14 | 0.25±0.02 | 0.19±0.01 |
| **30 mM** | Histamine+TTX | 1.19±0.16 | 6.49±0.61 | 0.32±0.01 | 1.09±0.04 |
| **15 mM** | Histamine+TTX | 1.09±0.12 | 3.41±0.54 | 0.27±0.02 | 0.63±0.05 |
| **5 mM** | Histamine+TTX | 0.46±0.03 | 0.91±0.07 | 0.26±0.01 | 0.17±0.01 |

**Figure 6.**
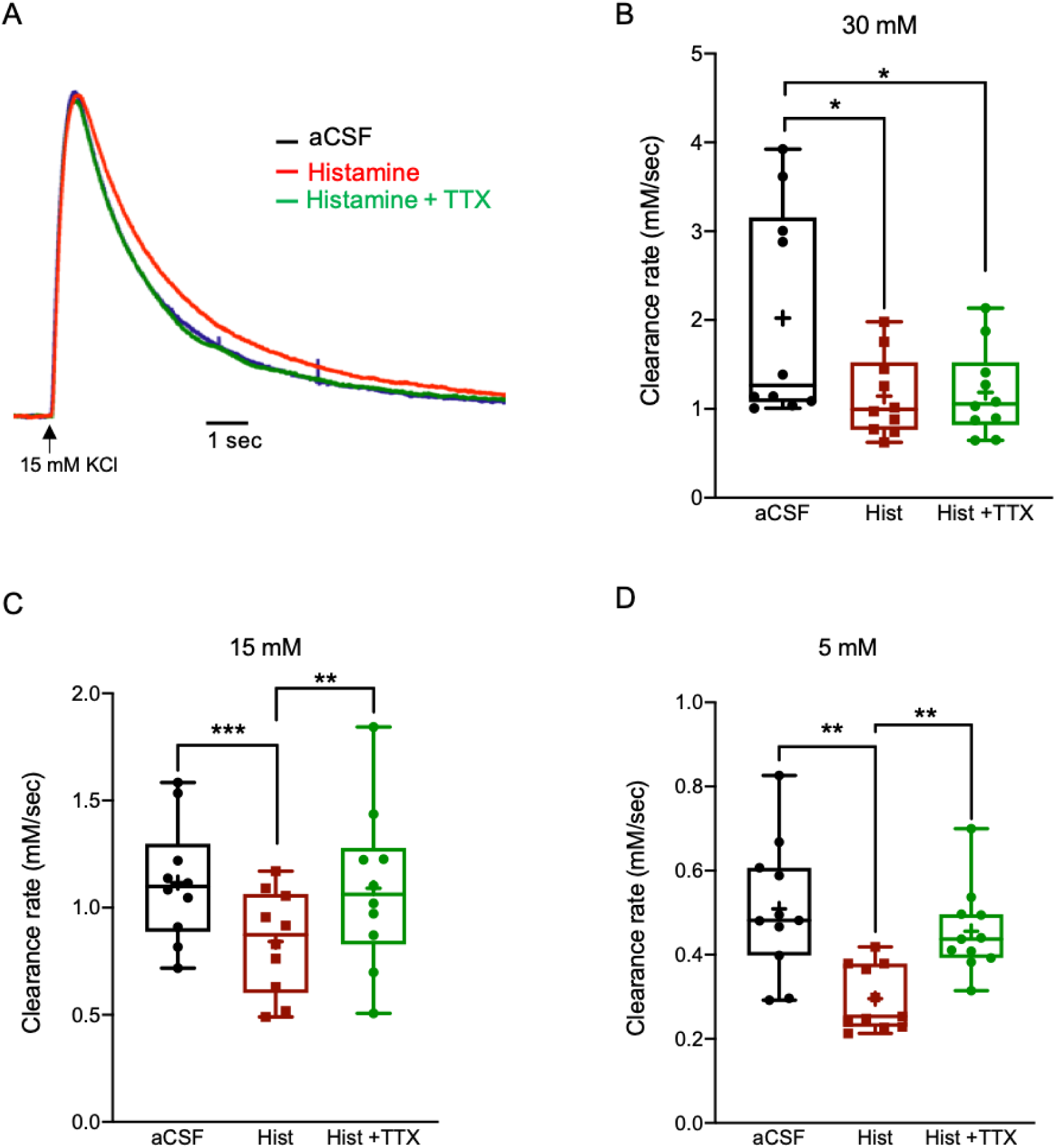
The impact of Histamine on the K^+^ clearance rate and astrocytic Ca^2+^ signalling. A) Sample traces of [K^+^]° recordings depicting the K^+^ clearance time course following local application of 15 mM KCl (arrow), before (aCSF, black) and after bath application of Histamine dihydrochloride (50 μM, red) or Histamine with 1 μM TTX (green). B-D) Box plots depicting the K^+^ clearance rate following local application of KCl at 30 mM (A), 15 mM (B) and 5 mM (C), before (aCSF, black dots) and after bath application of Histamine dihydrochloride (50 μM) without (red dots) or with TTX (green dots). Box plot definitions is the same as in Fig. 1. *p < 0.05; **p < 0.01; ***p < 0.0001

### Acetylcholine (ACh)

Astrocytes express both ionotropic receptors (α, β)(Oikawa *et al.*, 2005) and muscarinic G protein-coupled receptors (GPCRs) for ACh (M_1-3_)(Amar *et al.*, 2010; Hirase *et al.*, 2014). While activation of the Ca^2+^-permeable α7nACh receptor leads to [Ca^2+^]^i^ elevations due to Ca^2+^ entry from the extracellular mileu (Sharma & Vijayaraghavan, 2001), activation of M_1-3_ receptors in astrocytes increases [Ca^2+^]^i^ via activation of PLC, which elevates IP_3_ levels and promotes Ca^2+^ release from internal stores(Araque *et al.*, 2002),(Kelly *et al.*, 1996). Subsequently, astrocytic [Ca^2+^]^i^ elevations induce gliotransmitter release of glutamate, ATP or D-serine, thereby leading to modulation of synaptic strength and transmission in both the hippocampus (Papouin *et al.*, 2017) and the cortex (Takata *et al.*, 2011).

To test the impact of ACh on the K^+^ clearance rate, we bath applied slices with Carbachol (100 µM), a non-specific ACh agonist that binds and activates both nicotinic and muscarinic ACh receptors (Hobson *et al.*, 1983). However, the K^+^ clearance rate was comparable between normal aCSF and Carbachol conditions for all [K^+^]° tested, as shown in Figure 7A-D (30 mM KCl, 1.36±0.13 mM/sec; 15 mM KCl, 0.97±0.09 mM/sec; 5 mM KCl, 0.51±0.07 mM/sec; p > 0.05, One way ANOVA).

**Figure 7.**
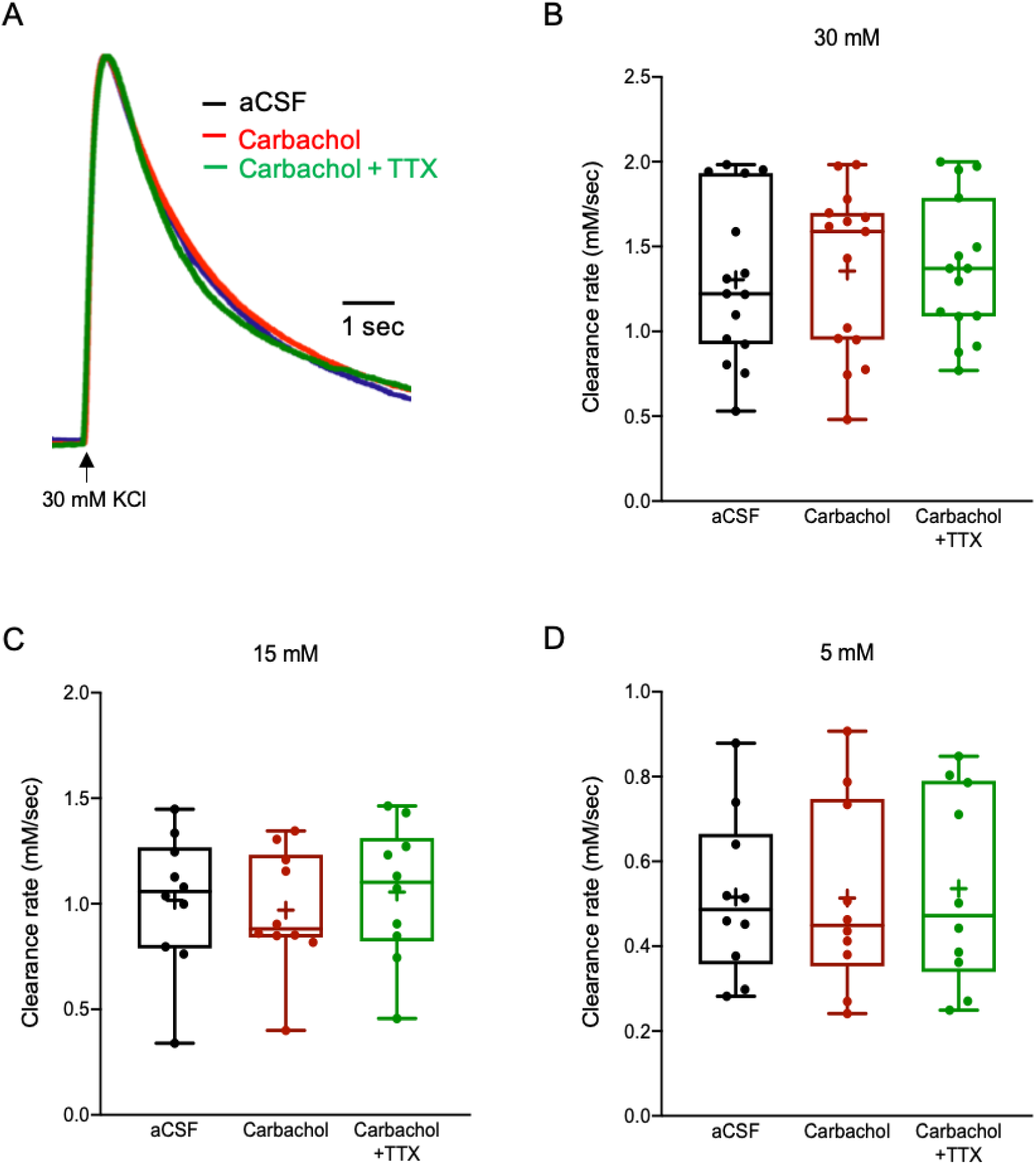
The impact of Carbachol on the K^+^ clearance rate and astrocytic Ca^2+^ signalling. A) Sample traces of [K^+^]° recordings depicting the K^+^ clearance time course following local application of 30 mM KCl (arrow), before (aCSF, blue) and after application of Carbachol (100 μM, red) or Carbachol with 1 μM TTX (green). B-D) Box plots depicting the K^+^ clearance rate following local application of KCl at 30 mM (B), 15 mM (C) and 5 mM (D), before (aCSF, black dots) and after bath application of Carbachol (100 μM) without (red dots) or with TTX (green dots). Box plot definitions is the same as in Fig. 1.

Since blockade of neuronal firing with TTX was comparable to control and Carbachol conditions (30 mM KCl, 1.37±0.11 mM/sec, n=15; 15 mM KCl, 1.06±0.10 mM/sec, n=10; 5 mM KCl, 0.54±0.07 mM/sec, n=10; p > 0.05, One way ANOVA, Figure 7A-C, Supplementary table 5), these results suggest that ACh has no direct or indirect impact on K^+^ clearance mechanisms.

**Table 5.** The impact of Acetylcholine on the K^+^ clearance rate. Average values of the [K^+^]° clearance rate (90-10%), amplitude, rise time (10-90%) and top peak area (10%) at all concentrations tested, before and after the application of Carbachol or Carbachol + TTX. Data is reported as mean ± S.E.M.

| [K<sup>+</sup>]<sub>o</sub> | Condition | Clearance<br>rate (mM/sec) | Amplitude<br>(mM) | Rise time<br>(sec) | Peak area<br>(mMxsec) |
| --- | --- | --- | --- | --- | --- |
| **30 mM** | aCSF | 1.30±0.12 | 6.63±0.4 | 0.31±0.02 | 1.29±0.07 |
| **15 mM** | aCSF | 1.02±0.10 | 3.26±0.4 | 0.27±0.02 | 0.83±0.09 |
| **5 mM** | aCSF | 0.52±0.06 | 1.34±0.2 | 0.22±0.01 | 0.26±0.03 |
| **30 mM** | Carbachol | 1.36±0.13 | 6.58±0.5 | 0.31±0.02 | 1.28±0.07 |
| **15 mM** | Carbachol | 0.97±0.09 | 3.28±0.4 | 0.27±0.01 | 0.77±0.07 |
| **5 mM** | Carbachol | 0.51±0.07 | 1.36±0.2 | 0.23±0.01 | 0.26±0.02 |
| **30 mM** | Carbachol+TTX | 1.37±0.11 | 6.79±0.6 | 0.30±0.02 | 1.22±0.08 |
| **15 mM** | Carbachol+TTX | 1.06±0.10 | 3.27±0.3 | 0.26±0.01 | 0.81±0.04 |
| **5 mM** | Carbachol+TTX | 0.54±0.07 | 1.39±0.2 | 0.23±0.01 | 0.25±0.01 |

## DISCUSSION

Neuronal activity is accompanied by a transient local increase in [K^+^]°, which must be cleared to maintain neuronal function. In the CNS, K^+^ homeostasis is maintained by astrocytic K^+^ clearance mechanisms, including “net K^+^ uptake” and K^+^ “spatial buffering” to distal areas through GJs (Orkand *et al.*, 1966), however the mechanisms that affect these clearance processes and overall [K^+^]° dynamics are largly unknown.

In this study, we investigated the impact of different neuromodulators known to act on both neurons and astrocytes in the K^+^ clearance process. Previous studies have demonstrated that neuromodulators, including DA (Ito & Schuman, 2007), ACh (Kirkwood *et al.*, 1999) and NA (O’Donnell *et al.*, 2012) affect neuronal excitability, leading to altered network oscillations at multiple frequencies (Constantinople & Bruno, 2011). Moreover, modulation of the cholinergic (Webster & Jones, 1988; Dort *et al.*, 2015; Ni *et al.*, 2016) or monoaminergic (Monti, 1993; Monti & Jantos, 2008; Carter *et al.*, 2010) signalling pathways has been reported to affect neural network oscillatory dynamics underlying behavioural shifts, as happens during different phases of sleep (i.e. REM *vs* NREM) or between sleep and arousal states. Another key modulator of extracellular K^+^ is the Na^+^/K^+^ ATPase (NKA pump), which is expressed in both neurons and astrocytes, though with different subunit isoforms (Larsen *et al.*, 2016). However, as there is no selective blockers for the astrocytic Na^+^/K^+^ ATPase, we did not measure its direct affect, to avoid misinterpertation of the direct impact of neuromodulators on astrocytic activity.

Recently Ding and colleagues showed that application of a cocktail of neuromodulators to cortical brain slices result in an increase of [K^+^]°, which did not involve neuronal activity (Kang *et al.*, 2016). Moreover, different behavioural states, such as arousal and sleep that are modulated by different neuromodulators, were found to be associated with alterations in [K^+^]° dynamics (Ding *et al.*, 2016). Consequently, we hypothesized that different neuromodulators can modulate [K^+^]° clearance rate by selectively activating different signalling pathways either directly (via astrocytes) or indirectly (via neurons) to increase [K^+^]° levels. To validate this hypothesis, we have measured the [K^+^]° clearance rate following local application of KCl at different concentrations in the presence of the neuromodulators 5-HT, DA, NE, Histamine and ACh (Carbachol).

### Mechanisms that affects the K^+^ clearance rate

Extracellular K^+^ dynamics are determined by the rate of active K^+^ uptake into nearby astrocytes, as well as the rate of extracellular diffusion (Gardner-Medwin, 1983). Previous reports suggested that the rate of K^+^ clearance can also be affected by different factors, including temperature (Ransom *et al.*, 2000*b*), ammonia (Rangroo Thrane *et al.*, 2013), glutamate (Enkvist & McCarthy, 1994) and pH, however, the cellular mechanisms affecting this clearance process are still largely unknown.

Among astrocytic K^+^ clearance mechanisms, K^+^ uptake becomes activated following low local increases in [K^+^]° (∼3-12 mM), mostly affecting small astrocytic networks located within close proximity to the synaptic release site, and becomes saturated at [K^+^]° above ceiling levels (>12 mM) (Heinemann & Dieter Lux, 1977; Orkand, 1986). In contrast, the K^+^ spatial buffering process via GJ-mediated astrocytic networks is active when there is high accumulation of [K^+^] (Bellot-Saez *et al.*, 2017). In that regard, agents that affect the clearance rate of low [K^+^]° (∼5 mM) independent of neuronal activity are likely to play a role in the modulation of astrocytic K^+^ uptake mechanisms, mediated via the NKA pump and Kir4.1 channels (Hajek *et al.*, 1996*a*; Butt & Kalsi, 2006; Larsen *et al.*, 2014), whereas compounds that affect the clearance rate of high and excessive [K^+^]° (15 mM and 30 mM respectively) are more prone to regulate the K^+^ spatial buffering process through GJs (Wallraff *et al.*, 2006; Pannasch *et al.*, 2011). Indeed, our results indicate that selective blockade of Kir4.1 channels affected the [K^+^]° clearance rate at all concentrations tested (Fig. 2), as the uptake occurs at all concentrations. However, selective inhibition of astrocytic GJ’s decrease the clearance rate only at high and excessive [K^+^]° (Fig. 2), consistant with previous reports (Wallraff *et al.*, 2006; Pannasch *et al.*, 2011). In addition, a key result in this study is that the rate of K^+^ clearance is concentration dependent and inversly correlated to the [K^+^]° (Fig. 1). This is probably due the fact that at high concentrations, K^+^ clearance is facilitated by GJ, as previously reported (Enkvist & McCarthy, 1994; Larsen *et al.*, 2014). A previous study specified that BaCl_2_ mainly affect the [K^+^]° peak amplitude (Larsen *et al.*, 2014), however the amplitude was measured following high frequency stimulation that lasted for 10 sec, during which neurons constantly excert K^+^ to the vicinity of the recording electrode. In comparrison, our stimulus was much shorter (0.1 sec), and therefore allow direct measurement of the clearance rate without the impact of further K^+^ increase to the extracellular fluid.

### Neuromodulators impact on the [K^+^]° clearance rate

The involvement of different neuromodulators in the regulation of network oscillations has been reported in many studies (Webster & Jones, 1988; Monti & Jantos, 2008; Dort *et al.*, 2015; Ni *et al.*, 2016), however the circuitry in which they mediate their impact remains elusive. Here we demonstrate that certain neuromodulators work in parallel on both neuronal and astrocytic networks, leading to differential impact on the K^+^ clearance rate. However, while some neuromodulators (e.g. 5-HT, DA and NA) exert their activity directly via astrocytes, other neuromodulators, such as Histamine expressed a more complex involvement, in which they affect the clearance rate indirectly via neuronal activity at low concentration and directly at excessive concentration (Fig. 6).

Moreover, while ACh had no impact on the [K^+^]° clearance rate at any of the concentrations tested (Fig. 7), all other neuromodulators significantly decrease the K^+^ clearance rate following excessive increase of [K^+^]° (Fig. 3-6). In contrast, the clearance rate following a low increase of [K^+^]° was affected only by DA and Histamine (Fig 4B and 6B respectively), though DA affected astrocytes direcly and the histaminergic effect was mediated by neuronal synaptic activity.

Together, these results suggest that 5-HT and NA effect on the K^+^ clearance rate was comparable to the impact of selective blockade of astrocytic gap junctions by Cx43 mimetic peptides, suggesting they affect only the spatial buffering process. In comparrison, DA and Histamine effect on the K^+^ clearance rate was comparable to the impact of selective blockade of Kir4.1 channels by BaCl_2_, suggesting they affect the uptake process. However, we cannot exclude the possibility that DA and Histamine affect the the spatial buffering process as well, as both processes are mediating K^+^ clearance under high and excessive concentrations.

Monoamines, including catecholamines (i.e. NA, DA), 5-HT and Histamine are involved in a broad spectrum of physiological functions (e.g. memory, emotion, arousal)(Cirelli & Tononi, 2000; Lowry *et al.*, 2005; Eckart *et al.*, 2016), as well as in psychiatric and neurodegenerative disorders (e.g. Parkinson’s disease, Alzheimer’s disease, schizophrenia, depression)(Panula *et al.*, 1997; Ray *et al.*, 2008; Heneka *et al.*, 2010). At the cellular level, synaptic release of neuromodulators impact membrane properties as well as intracellular signalling pathways in both neurons and glial cells (Kang *et al.*, 2016; Ma *et al.*, 2016), and previous reports showed that different neuromodulators can fine-tune the hyperpolarization-activated current *I*_*h*_ (Maccaferri & McBain, 1996; Rosenkranz, 2006; Ma *et al.*, 2007), thereby affecting membrane resonance of individual neurons, which affect the oscillatory behaviour of single neurons and their synchronization into networks (Tseng *et al.*, 2014). However, whether this was a direct effect of the neuromodulators on neuronal activity, or indirect via astrocytic modulation was never tested.

A recent study by Ma and colleagues show that neuromodulatory signalling in Drosophila can flow through astrocytes and promote their synchrous activation (Ma *et al.*, 2016). They further suggest that astrocyte-based neuromodulation is an ancient feature of the Metazoan nervous sytem. Moreover, Slezak and colleagues (Slezak *et al.*, 2018) suggested that astrocytes function as multi-modal integrators, encoding visual signals in conjuction with arousal state. Our results support this concept, in which neuromodulators impact network oscillatory activity through parallal activation of both neurons and astrocytes, establishing a synergetic mechanism to maximise their impact on synchronous network activity and recruitment of neurons into networks.

## ADDITIONAL INFORMATION

### Author Contributions Statement

YB conceived the project. YB, JWM, YBA and ABS designed the experiments. ABS and OK performed and analysed the electrophysiological recordings. All authors wrote and approved the paper.

## Acknowledgments

We thank Dr. Sindy Kueh for technical assistance. This study was supported by IPRA to ABS and the Ainsworth medical research innovation fund awarded to YB and JWM.

